# Nitrogen starvation causes lipid remodeling in *Rhodotorula toruloides*

**DOI:** 10.1101/2023.05.07.539759

**Authors:** Shekhar Mishra, Anshu Deewan, Huimin Zhao, Christopher V. Rao

## Abstract

The oleaginous yeast *Rhodotorula toruloides* is a promising chassis organism for the biomanufacturing of value-added bioproducts. It can accumulate lipids at a high fraction of biomass. However, metabolic engineering efforts in this organism have progressed at a slower pace than those in more extensively studied yeasts. Few studies have investigated the lipid accumulation phenotype exhibited by *R. toruloides* under nitrogen limitation conditions. Consequently, there have been only a few studies exploiting the lipid metabolism for higher product titers. Here, we present a multi-omic investigation of the lipid accumulation phenotype under nitrogen limitation. Through an integrative lens of transcriptomic and lipidomic analysis, we identify that *R. toruloides* undergoes lipid remodeling during nitrogen limitation, wherein the pool of phospholipids gets remodeled to mostly storage lipids. This insight into the mechanisms of lipid accumulation can lead to the success of future metabolic engineering strategies for overproduction of oleochemicals.

**Highlights:**

- The oleaginous yeast *R. toruloides* displays enhanced lipid accumulation during nitrogen starvation.
- A multi-omic investigation of the lipid accumulation phenotype was carried out.
- Lipid remodeling was observed during the accumulation phase, wherein carbon was transferred from phospholipids to storage lipids.
- Multi-omic analysis suggested that selective regulation within lipid biosynthesis controls for the specific increase of storage lipids.

## 1. Introduction

The unicellular microorganism yeast is a promising platform for industrial-scale fermentations of value-added compounds. As a model yeast, *Saccharomyces cerevisiae* has been extensively characterized and engineered to produce a wide variety of bioproducts (Lian et al., 2018; Volk et al., 2022). However, *S. cerevisiae* is not an ideal host for overproducing lipids and lipid-related chemicals, also called oleochemicals. Oleaginous yeasts such as *Yarrowia lipolytica* and *Rhodotorula toruloides*, which can accumulate a much higher fraction of their biomass as lipids, are theoretically better platforms for producing oleochemicals (Beopoulos et al., 2009; Wen et al., 2020). The basal metabolism of these yeasts results in a higher yield of lipids per substrate consumed which is required to make such industrial fermentations economically viable. The red yeast *R. toruloides* is capable of accumulating as high as 70% of its biomass as lipids, while also allowing high cell density cultures, thus resulting in high titers of lipid production (Lopes et al., 2020; Zhang et al., 2016a). Besides a high flux in lipid biosynthesis, *R. toruloides* is also known to show high tolerance towards inhibitory compounds, specifically those found in lignocellulosic biomass hydrolysate (Sundstrom et al., 2018; Yaegashi et al., 2017; Zhao et al., 2012). This tolerance combined with an ability to consume hexose and pentose sugars, both found in lignocellulosic biomass hydrolysate, suggests that *R. toruloides* can serve as a better host for converting lignocellulosic sugars into value-added compounds. As *R. toruloides* accumulates high levels of lipids and carotenoids, both of which are synthesized from acetyl-CoA, *R. toruloides* has the potential to serve as a platform strain for producing a variety of compounds synthesized from acetyl-CoA. Previous studies have explored overproducing lipids in *R. toruloides* through a variety of strategies like inhibiting fatty acid degradation, increasing the flux of acetyl-CoA or other biosynthesis reactions in lipid pathways (Zhang et al., 2016a, 2016b). Other work has focused on the production of fatty alcohols and triacetic acid lactone, both of which are synthesized from acetyl-CoA (Cao et al., n.d.; Schultz et al., 2022). The strategies implemented in most metabolic engineering studies so far focus on similar strategies of reducing lipid catabolism, increasing precursor supply, and introducing a heterologous enzyme (push-pull-block strategy).

While *R. toruloides* appears to be a promising host for the biomanufacturing of oleochemicals, the microorganism poses several challenges. It is considered a non-model yeast and as such, its metabolism is not well understood. While the accumulation of lipids in low nitrogen conditions has been well reported, the molecular basis of this phenotype has not been clearly elucidated (Zhu et al., 2012). Thus, to harness the full potential of *R. toruloides* in metabolic engineering applications, it is desirable to understand the mechanism of lipid accumulation.

Within systems biology, transcriptomic analysis is a robust analytical technique providing insights into the transcriptional machinery of an organism. Other studies endeavoring to find the molecular basis of lipid accumulation in *R. toruloides* have also employed transcriptomic analysis for their investigations. Coradetti et al. performed transcriptomic analysis of a library of *R. toruloides* loss-of-function mutants to identify putative genes affecting lipid metabolism in the yeast (Coradetti et al., 2018). Zhu et al. employed a multi-omic method where genomic, transcriptomic and proteomic data were integrated to identify correlations between lipid accumulation and nitrogen compound recycling (Zhu et al., 2012). Recently, Jagtap et al. performed a transcriptomic and metabolomic analysis of *R. toruloides* grown on different sugars, identifying regulation patterns in central metabolic pathways as a result of growth on different substrates (Jagtap et al., 2021). In other yeasts such as *Y. lipolytica*, multi-omic analysis, which included a genome-scale model, metabolite profiling, lipidomic analysis and transcriptomic data from RNA-Seq, identified the origin of lipid accumulation in flux rewiring within the amino acid metabolism (Kerkhoven et al., 2016). However, in *R. toruloides*, no such integrated transcriptomic and lipidomic analysis has been performed so far.

In this study, we undertake a multi-omic investigation of lipid accumulation in *R. toruloides*. Nitrogen-sufficient and nitrogen-deficient conditions were studied to contrast a neutral phenotype against a lipid accumulation phenotype. The cells grown in both conditions were harvested for both transcriptomic and lipidomic analyses. As a primary analysis, clustering analysis of the transcriptomic data was qualitatively correlated against a similar trend in lipidomic data. Hierarchical clustering of the lipid data, however, showed a different clustering pattern than that seen from RNA-Seq clustering. Multi-omic analysis of the transcriptomic and lipidomic data indicated a lipid remodeling event wherein glycerophospholipids pathways are repressed whereas most of the carbon is shifted towards storage lipids. Our work represents the first comprehensive study integrating transcriptomic and lipidomic analyses of lipid accumulation phenotypes in *R. toruloides*.

## 2. Materials and methods

### 2.1 Strains, media, and growth construction

*R. toruloides* IFO0880, mating type A2, was obtained from the NITE Biological Resource Center in Japan (NBRC 0880). YPD medium (10 g/L yeast extract, 20 g/L peptone, and 20 g/L glucose) was used for growth of *R. toruloides* precultures. A single colony from a YPD agar plate was inoculated into 2 mL of YPD liquid medium to obtain *R. toruloides* seed cultures. Seed cultures were then used to inoculate 25 mL of media with a defined carbon-to-nitrogen ratio in a 125-mL baffled shake flask with a starting OD_600_ of 1. The optical density at 600 nm or OD_600_ was used to monitor the cell density in liquid cultures. OD_600_ of 1.0 corresponds to roughly 10^7^ cells per mL. The cells were then grown at 30 °C and 250 rpm. The formulation of media for different C/N ratios is listed in Table S1.

### 2.2 Sample extraction for RNA-seq

Seed cultures at exponential phase were collected and centrifuged at 6000 × g for 3 min at 4°C. Supernatant was discarded and the pellets were resuspended in 1 mL of ddH_2_ O. Seed cultures then used to inoculate 25 mL of growth media with defined carbon-nitrogen ratios in a 125-mL baffled shake flask with a starting OD_600_ of 1 and incubated at 30 °C and 250 rpm. Growth experiments are performed with three biological replicates. Samples were collected at defined timepoints as described in Section 3.1.

### 2.3 Experimental procedure for RNA-Seq

The cell cultures containing a total OD of 30 were collected in centrifuge tubes and centrifuged at 6000 × g for 3 min at 4 °C. Supernatant was discarded and pellet was used for RNA extraction. Total RNA was extracted using the RNeasy mini kit (Qiagen, Hilden, Germany) as previously described, with a slight modification (Jagtap et al., 2019; Zhang et al., 2019). The

*R. toruloides* cell pellet was resuspended in 350 μL of Buffer RLT from the RNeasy mini kit (Qiagen, Hilden, Germany). Approximately 500 μL of acid-washed glass beads (acid washed, 425–600 μm; Sigma, St. Louis, MO, USA) was added and homogenized using a FastPrep-24 homogenizer (MP Biomedicals, Irvine, CA, USA), beaten at a speed of 5 m/s for 30 s six times with cooling on ice between beatings. The cell lysates were purified according to the kit’s protocol titled “purification of total RNA from yeast.” Extracted RNA was then treated with Turbo RNase-free DNase kit (ThermoFisher, Waltham, MA, USA) according to the manual and purified again with the RNeasy mini kit protocol titled “RNA clean up.” The stranded RNAseq libraries were prepared with Illumina’s TruSeq Stranded mRNA Sample Prep kit. The libraries were quantitated by qPCR and sequenced on one lane for 101 cycles from one end of the fragments on a HiSeq 4000 (Illumina, San Diego, CA, USA). Fastq files with 100 bp reads were generated and demultiplexed with the bcl2fastq v2.17.1.14 Conversion Software (Illumina, San Diego, CA, USA).

### 2.4 RNA-Seq data analysis

To obtain gene expression profiles during growth of *R. toruloides* IFO0880 on different substrates, total RNA was extracted, and a mRNA focused library was sequenced. Adaptor sequences and low-quality reads were trimmed using Trimmomatic (Bolger et al., 2014). Trimmed reads were analyzed for quality scores using FastQC (“Babraham Bioinformatics - FastQC A Quality Control tool for High Throughput Sequence Data,” n.d.). Reads were mapped to the *R. toruloides* IFO0880 v4.0 reference genome (NCBI Accession GCA_000988875.2) with STAR version 2.5.4a (Coradetti et al., 2018; Dobin et al., 2013). Between 95 and 98% of the reads were successfully mapped to the genome for each sample. Read counts were calculated using featureCounts from the Subread package, v1.5.2 (Liao et al., 2014). Differential expression analysis was performed on the reads counts in R v4.0.5 (Cite R core team) using edgeR v3.32.1 and limma v3.46.0 (Ritchie et al., 2015; Robinson et al., 2010). Graphical representation of expression data was constructed using R packages: PCAtools v2.2.0, gplots v3.1.1, and Glimma v2.0.0 (Blighe et al., 2022; Su et al., 2017; Warnes et al., 2005). Before plotting heatmaps, the data was normalized row-wise (using the scale function in R), first by centering (subtracting the row mean from each value) and then scaling (dividing each data point by row’s standard deviation). Heatmaps were plotted using heatmap.2 function from gplots. Genome sequence, gene models, and functional annotation of *R. toruloides* was downloaded from the DOE Joint Genome Institute’s Mycocosm portal (Coradetti et al., 2018; Grigoriev et al., 2014).

### 2.5 Sample preparation for lipidomic analysis

At the time of sampling, cell concentration of the culture was measured as OD_600_ /mL. One mL of cell culture was harvested, washed once with 150 mM ammonium bicarbonate (ABC) buffer (pH = 8), then resuspended in 500 μL ABC, and lysed open with approximately 200 μL of 0.5μm zirconium glass beads via high-speed vortexing for 30 minutes. For lipid extraction, 2 OD_600_ units of cell lysate were added to a glass tube containing 200 μL ABC, 1 mL of 2:1 chloroform:methanol and 12 μL of an internal standard cocktail (mole amounts of each lipid in the cocktail listed in Table S2). The glass tube was vortexed in a Fisherbrand MultiTube Vortexer (Thermo Fisher Scientific, Waltham, MA) for 2 hours at 2500 rpm. After the phases were clearly separated, the chloroform layer was separated into a fresh vial and dried overnight. The dried lipid extract was resuspended in 100 μL of 4:2:1 isopropanol:methanol:chloroform. 10 μL of the lipid extract was injected on an LC-MS instrument (Vanquish UHPLC and Q-Exactive Orbitrap, Thermo Fisher Scientific, Waltham, MA).

### 2.6 Lipidomic data generation and data analysis

The LC-MS protocol was followed as described in (Zhang et al., 2018). LC separation was performed on a Thermo Accucore Vanquish C18+ column (2.1 × 150 mm, 1.5 μm) with mobile phase A (60% acetonitrile: 40% H2O with 10 mM ammonium formate and 0.1% formic acid) and mobile phase B (90% isopropanol: 10% acetonitrile with 10 mM ammonium formate and 0.1% formic acid) and a flow rate of 0.25 mL/min. The linear gradient was as follows: 0 min, 60% A; 12 – 13.5 min, 0% A; 14 – 16 min, 60% A. The gas flow rates and MS1/MS2 scan parameters were followed exactly as listed in (Zhang et al., 2018). Data processing of the .RAW files generated from LC-MS runs was performed using the MS-DIAL software (Tsugawa et al., 2015). Identity of lipids was ascertained by comparing spectra to an in-house database of lipid molecules. Finally, quantification was performed using a one-point calibration where the peak intensity of each lipid molecule was normalized to the intensity of the representative lipid for its class within the spiked-in internal standard cocktail. This normalized value was multiplied to the absolute mole amount of the internal standard.

## 3. Results

### 3.1 Experimental design for studying the lipid accumulation phenotype

The growth and lipid accumulation phenotypes of *R. toruloides* IFO0880 were studied in three different media conditions. The three conditions were formulated to contain C/N ratios of 5, 100 and 150. A C/N ratio of 5 represented a baseline media with nitrogen in sufficient concentration, whereas media with C/N ratios of 100 and 150 were nitrogen-deficient and known to promote lipid accumulation. Samples were collected at three timepoints during growth: 8 hours (early exponential phase), 12 hours (mid-exponential phase) and 36 hours (early saturation phase). For the two nitrogen-deficient conditions (C/N of 100 and 150), an additional sample at 88 hours of growth was added (Figure 1).

**Figure 1.**
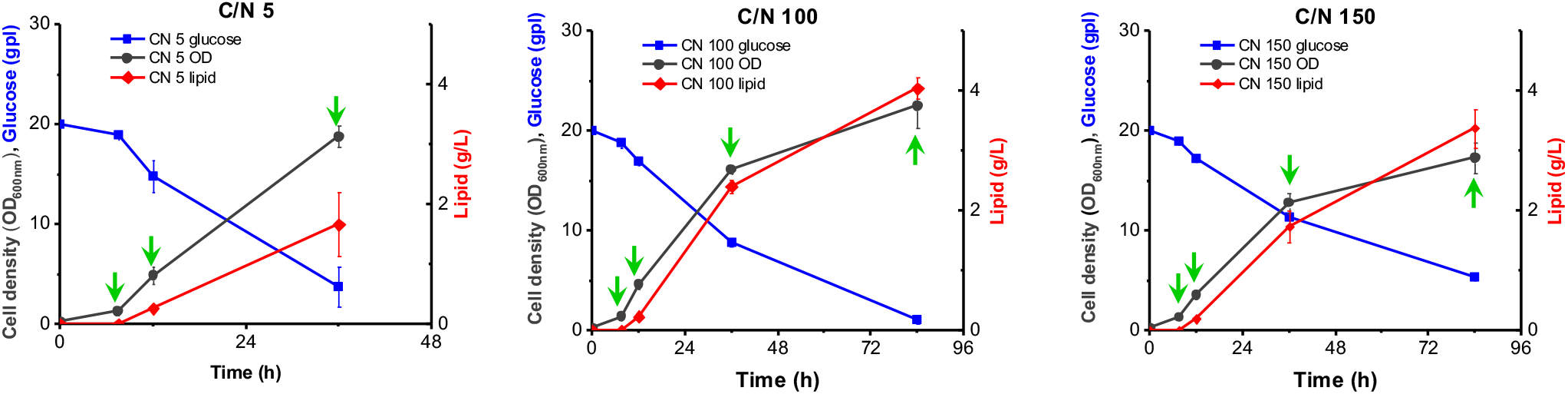
Experimental design scheme depicting the growth curves of *R. toruloides* IFO0880 in media with different C/N ratios. The green arrows in each plot represent the times when culture samples were collected for transcriptomic and lipidomic analysis.

### 3.2 Transcriptomic analysis of lipid accumulation dynamics

RNA-Seq analysis was used to inspect the differential gene expression in *R. toruloides* IFO0880 across different conditions and timepoints as described above. Hierarchical clustering of the RNA-Seq data showed that during the early exponential growth phase, when nitrogen is still available in all three media conditions, the transcriptomes of cells grown in the different media conditions cluster together (Figure 2). As nitrogen depletion took effect during mid-exponential phase, the cultures grown in C/N of 100 and 150 began to diverge from C/N of 5, where nitrogen was still expected to be available. This divergence is highlighted even further in the early and late-saturation samples where nitrogen-depleted conditions cluster together.

**Figure 2.**
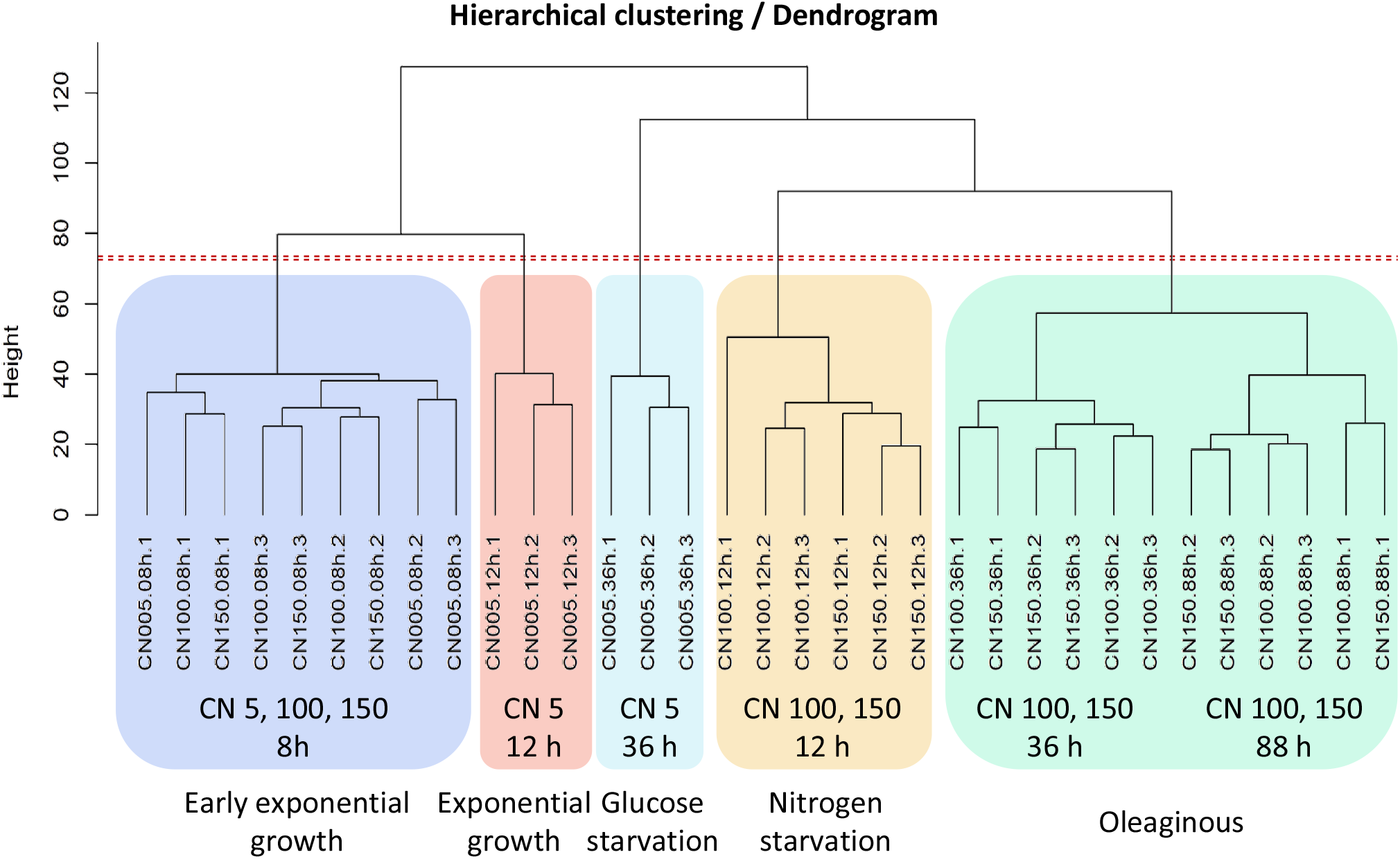
Hierarchical clustering performed on the time-based RNA-Seq analysis of *R. toruloides* cultures. The samples are shaded using different color boxes to lump together samples from different phases of growth. From left to right: Early exponential, Exponential, Glucose starvation, Nitrogen starvation and Oleaginous growth phases.

Principal component analysis (PCA) aided in improved visualization and analysis of this divergence where the lipid-accumulating and non-lipid-accumulating samples clearly follow different trajectories and cluster separately (Figure 3).

**Figure 3.**
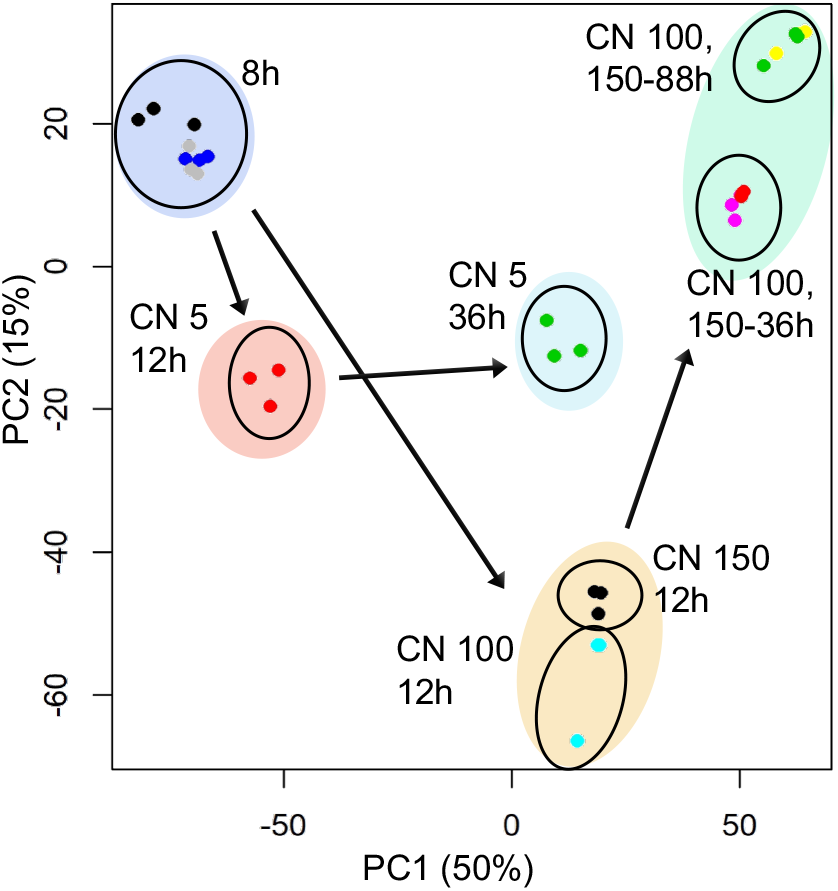
Principal component analysis of the RNA-Seq dataset showing the clustering of nitrogen-sufficient and nitrogen-deficient cultures. Additionally, cultures also cluster together based on the growth phase, with a clear divergence emerging at the exponential growth phase cultures (12 hours of growth).

RNA-Seq data was used to identify the subset of genes that were differentially expressed in conditions of nutrient starvation compared to baseline cultures. A threshold of greater than 2-fold change in gene expression (which includes up- and down-regulation) was employed. From the identified differentially expressed genes, the fraction of up- and down-regulated genes in all annotated pathway classes for *R. toruloides* IFO0880 were calculated and displayed as a horizontal bar graph. Figure 4 shows one such analysis for the comparison of differentially expressed genes between C/N of 100 and 150 compared to C/N of 5 at the 36-hour timepoint. Since the same set of genes were differentially expressed in both C/N 100 and 150 in comparison with C/N 5, the conditions could be plotted interchangeably. The figure shows the metabolic classes with the highest fraction of up-regulated genes at the top and those with the highest fraction of down-regulated genes at the bottom. Inspection of this plot shows that during nitrogen starvation in C/N 100 and 150, the metabolic processes of translation, transcription and secondary metabolism were entirely down-regulated. Genes involved with amino acid metabolism also see a major fraction of differential expression as down-regulation.

**Figure 4.**
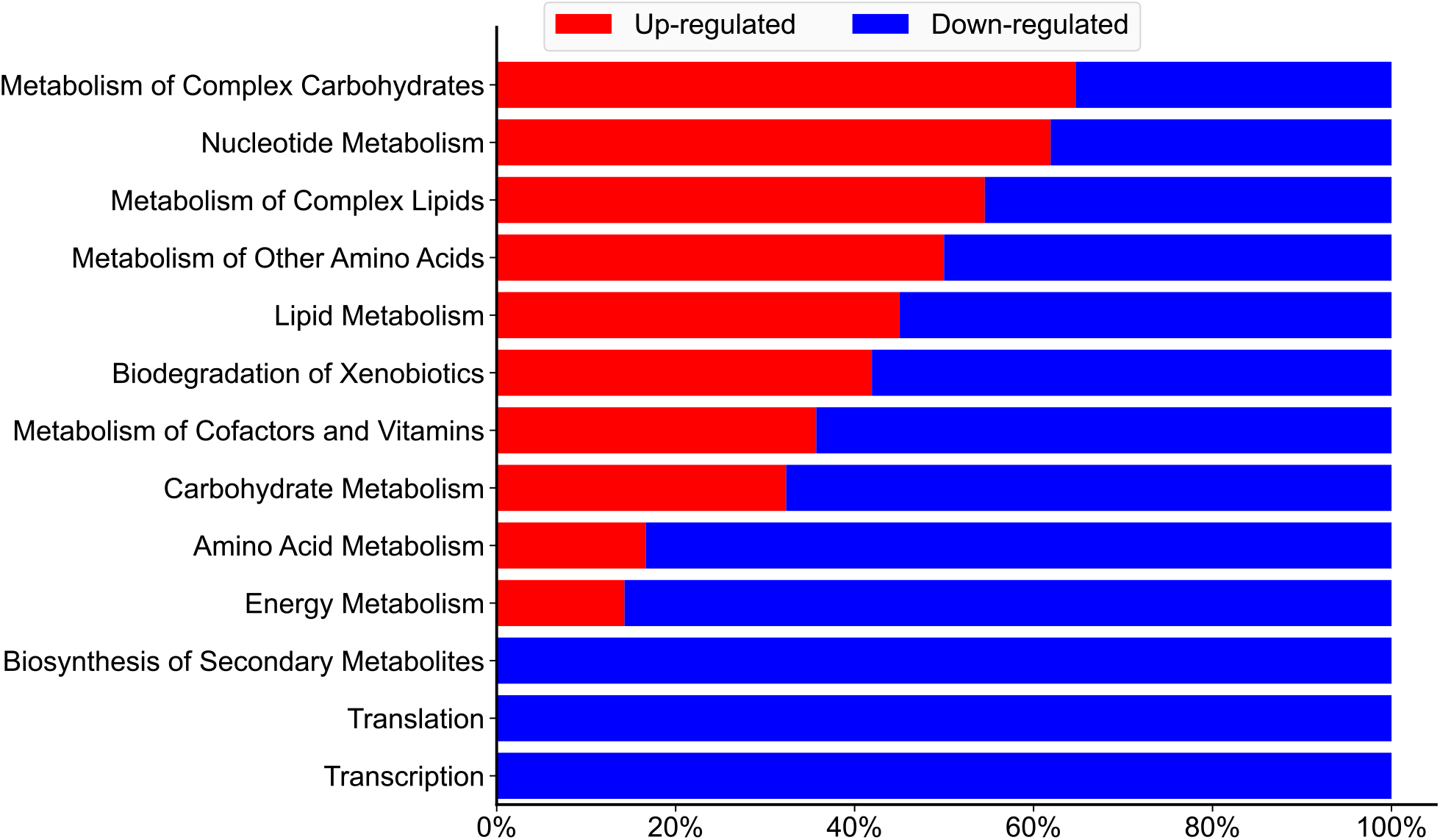
Gene set analysis of nitrogen limitation in *R. toruloides* IFO0880. The differentially expressed genes were sampled at a timepoint of 36 hours and contrasted between nutrient limited conditions of C/N 100 and 150 versus C/N 5 (which served as a baseline). Since the same set of genes were differentially expressed in both C/N 100 and 150 in comparison with C/N 5, the conditions could be plotted interchangeably.

This trend has also been noticed in *Y. lipolytica* where nitrogen starvation significantly down-regulated amino acid metabolism (Kerkhoven et al., 2016). However, in contrast with the findings of the *Y. lipolytica* study, genes involved in lipid metabolism were more differentially regulated in *R. toruloides* compared to the baseline condition. The study in *Y. lipolytica* reported minimal regulatory impact of nitrogen starvation on lipid metabolic genes, whereas our data indicated significant regulation in *R. toruloides*.

### 3.3 Lipidomic analysis of lipid accumulation dynamics

Liquid chromatography – mass spectrometry (LC-MS) was used to perform lipidomic analysis of extracted lipids from *R. toruloides* IFO0880 cells. The cells were harvested at the timepoints denoted by green arrows in Figure 1. This allowed a time-based inspection of the lipidome for both lipid-accumulating and non-lipid-accumulating phenotypes.

At the earlier timepoints, the measured lipid concentrations followed a similar trend to the expected clustering behavior observed from RNA-Seq data shown in Figure 2 and Figure 3. Lipid concentrations for the early exponential samples (8-hour growth) showed similar levels for all major lipid classes (Figure 5). The divergence of lipid profiles at the exponential (12-hour sample) and saturation phase (36-hour sample) was noticeable in the phospholipid pools (Figure 5 A-D). Most glycerophospholipids (phosphatidylcholine (PC), phosphatidylethanolamine (PE), phosphatidylinositol (PI) and phosphatidylglycerol (PG)) were higher in C/N of 5 compared to C/N of 100 or 150 (Figure 5 A-D).

**Figure 5.**
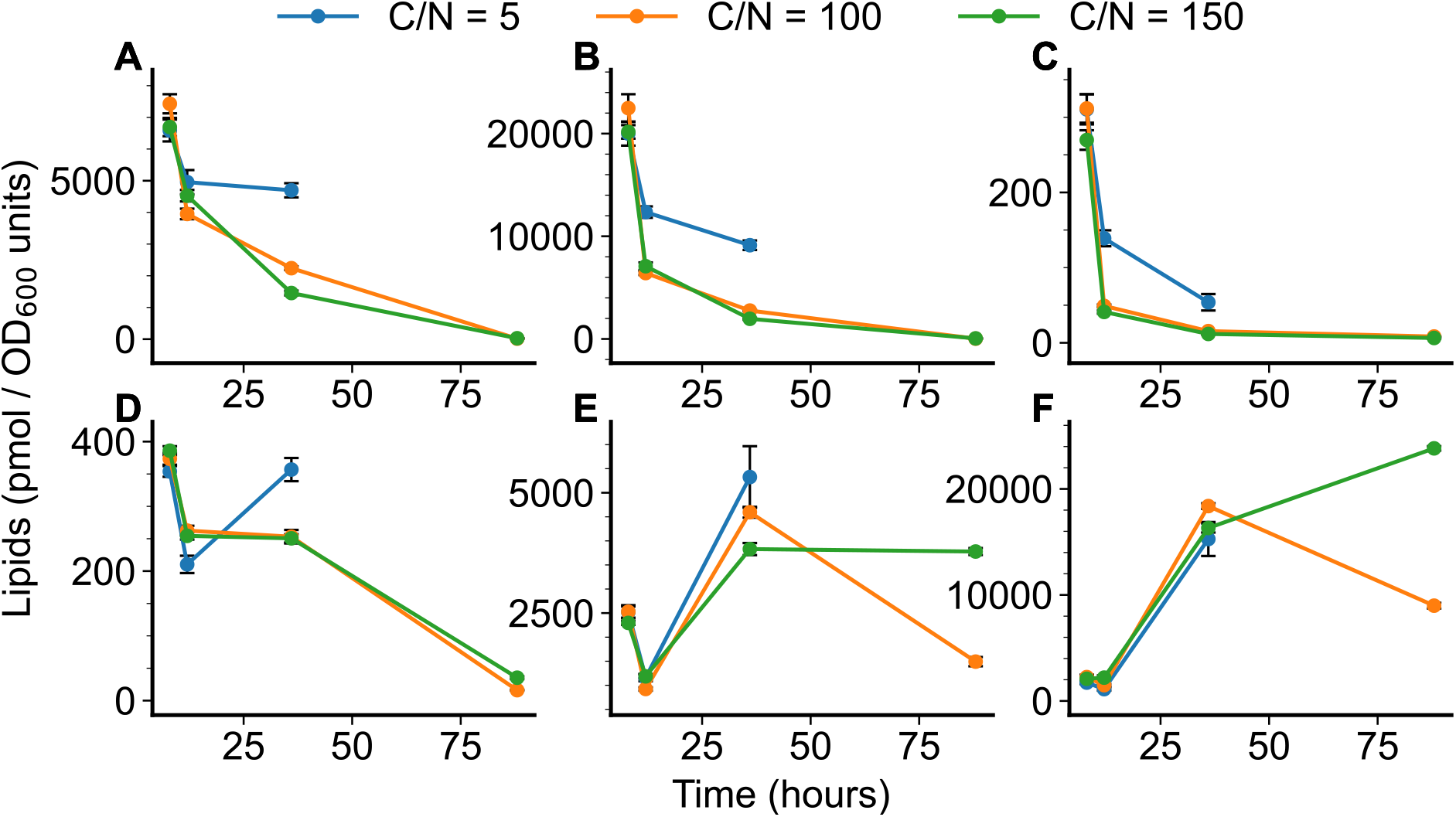
Lipidomic analysis of *R. toruloides* IFO0880 strain cultivated in different growth media containing C/N ratios of 5, 100 and 150, and sampled at various timepoints. Shown here are quantified lipid classes of all three growth conditions sampled at 8 hours, 12 hours, 36 hours and 88 hours of growth. A) Phosphatidylcholine (PC), B) Phosphatidylethanolamine (PE), C) Phosphatidylglycerol (PG), D) Phosphatidylinositol (PI), E) Diacylglycerol (DAG), and F) Triacylglycerol (TAG).

At the late saturation phase of oleaginous lipid accumulation, samples were only collected for C/N of 100 and 150, and hence, lipid measurements were only available for lipid accumulating phenotypes. At this late stage of growth, the fraction of lipids within the cells is mostly TAG. Analysis of TAG levels within the two lipid-accumulating phenotypes shows a divergence in TAG levels with C/N 150 continuing to store carbon as TAG even after 88 hours of growth, while TAG levels show a decrease in C/N 100 at the same timepoint (Figure 5F). Extracellular measurements (Figure 1) show that C/N 100 has consumed most of the glucose by this point in growth while significant amounts of glucose remain unconsumed in C/N 150. This measurement combined with the lipidomic dataset suggests that in the C/N 150 media, *R. toruloides* continues to accumulate carbon into TAG much longer, while C/N 100, having depleted most of its sugar substrate, potentially relies on lipid remodeling to generate carbon for continued survival. Within the set of annotated mRNA transcripts that were differentially expressed, the glycerol dehydrogenase was up-regulated in the C/N 100 condition at 88 hours compared to the C/N 150 sample, suggesting increased activity in glycerolipid catabolism.

Further inspection of the distribution of lipids showed that in all three growth conditions, the distribution of lipids shifts from mostly phospholipids towards mostly storage lipids. At the start of each growth culture (before nitrogen starvation takes effect), phospholipids dominate the fraction of intracellular lipids. As each growth condition progresses with time, the fraction of phospholipids was seen to subside and replaced almost entirely by storage lipids (Figure 6). While culture conditions of C/N 100 and 150 closely resembled each other in trajectory, even C/N 5 showed a similar trend, albeit with different values.

**Figure 6.**
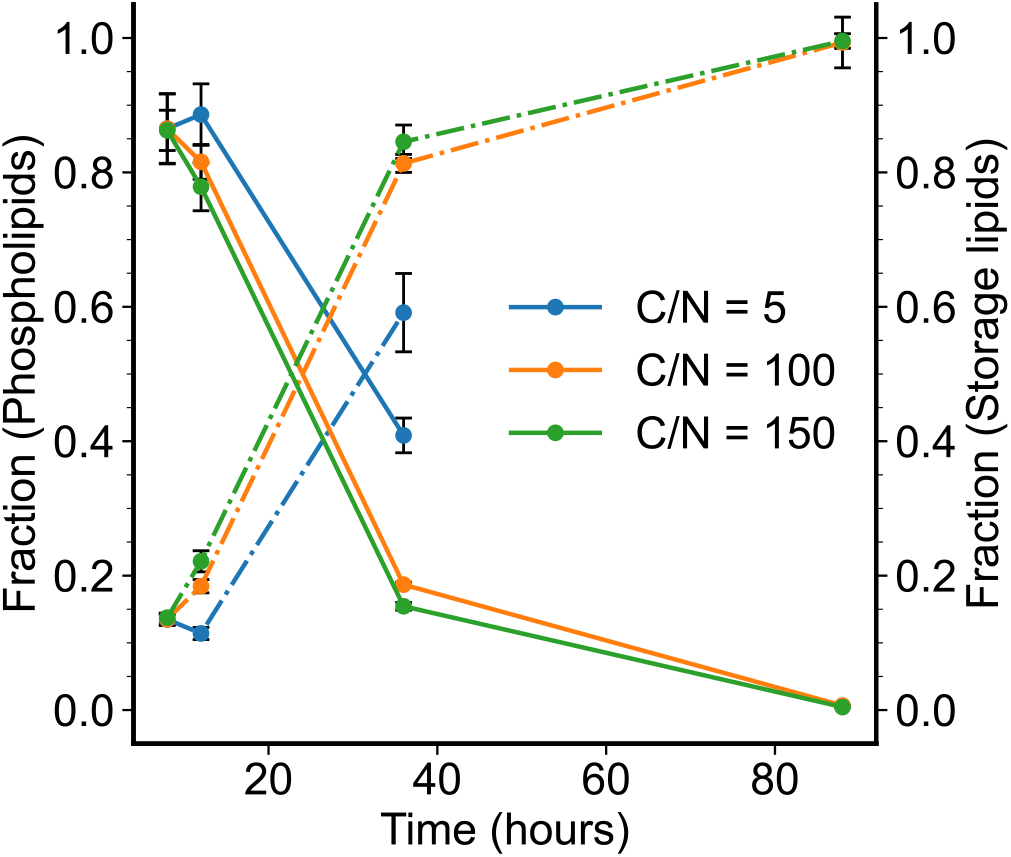
Variation in the fraction of total lipid pool in *R. toruloides* IFO0880 at each time point and for each growth condition. Two fractions are calculated: fraction of phospholipids (solid line) and fraction of storage lipids (dash-dot line).

A shift in the total intracellular lipids from phospholipids to storage lipids further reinforces the notion of an active lipid remodeling process as nitrogen starvation starts to take effect. Studies in the model oleaginous yeast, *Y. lipolytica*, have demonstrated that nitrogen limitation causes an overall increase in lipid pools in that organism (Kerkhoven et al., 2016; Morin et al., 2011). They show that nitrogen limitation in *Y. lipolytica* causes little change to transcriptional regulation in lipid metabolism and instead modulates amino acid metabolism. The overall increase in lipid levels has been attributed to an overflow of carbon into lipid biosynthetic pathways (Kerkhoven et al., 2016). In contrast, however, the results shown in Figure 4 and Figure 6 suggest that nitrogen starvation in *R. toruloides* is accompanied by significant lipid regulation.

A hierarchical clustering analysis of the lipidomic data was performed (Figure 7), which shows that the 88-hour and 36-hour time points cluster similarly while the 8-hour and 12-hour time points cluster together. The latter suggests that the divergence in the RNA-Seq data that appears at 12 hours of growth manifests in lipid phenotypes with a time delay, which is generally observed in RNA-metabolite trends (Takahashi et al., 2011).

**Figure 7.**
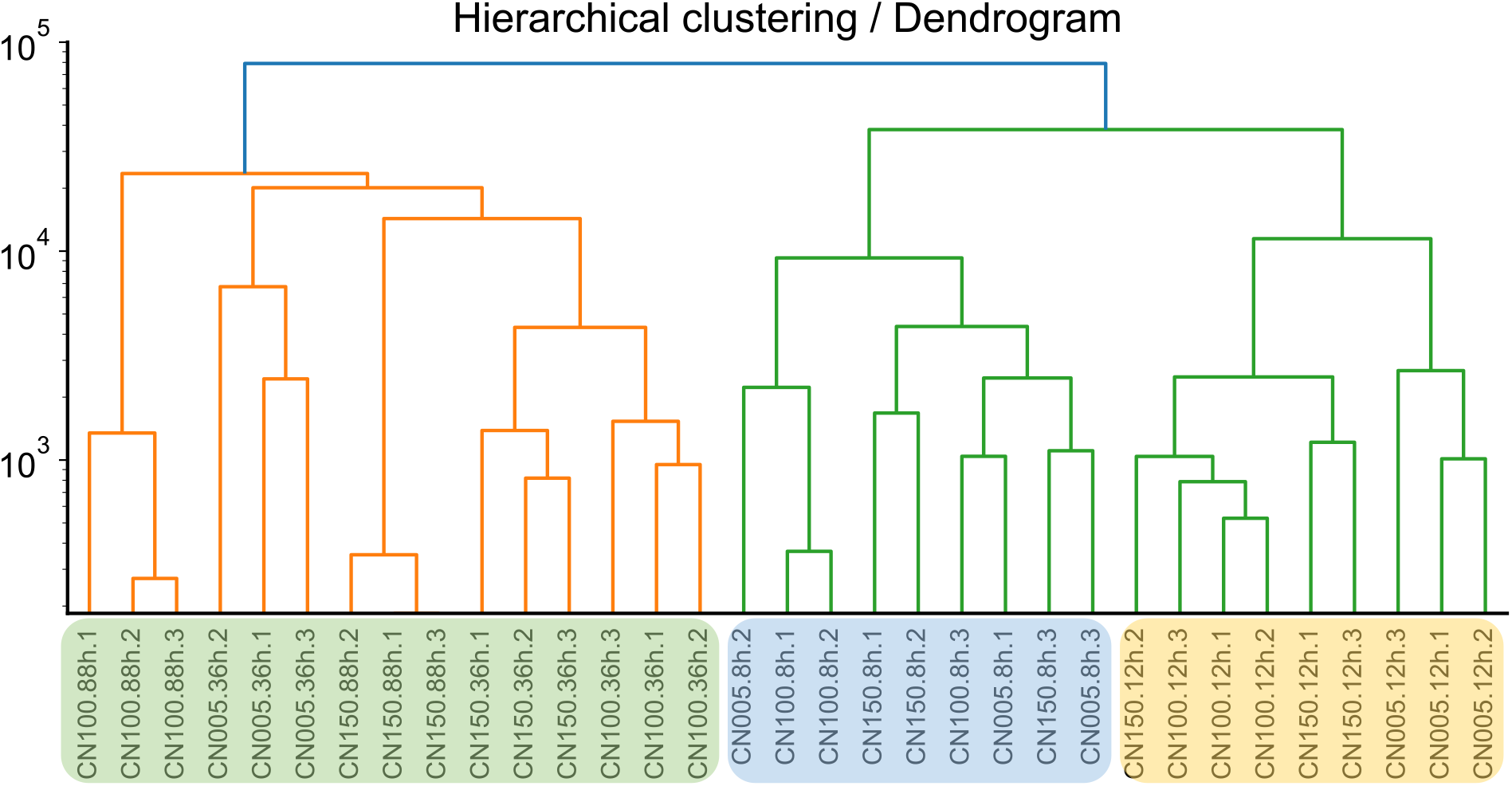
Hierarchical clustering performed on the time-based lipidomic analysis of *R. toruloides* cultures.

### 3.4 Multi-omic integrated analysis of lipid accumulation

So far, all literature reported investigations on the lipid accumulation phenotype of *R. toruloides* IFO0880 were performed using individual omic analyses, namely transcriptomic and lipidomic analysis in isolation. However, each of these datasets only present a partial picture of the biological processes governing the oleaginous yeast’s behavior. The transcriptomic dataset indicates snapshots of the gene expression profile of the cells, whereas the lipidomic dataset conveys a snapshot of the lipidome. Thus, a systematic method of analysis that reasonably integrates both datasets was sought for analysis.

An integrative method of analysis previously described in Hsu and coworkers (Hsu et al., 2017) was employed wherein transcriptomic and metabolomic profiles were analyzed together in the form of a cluster map. The 4 quadrants of such a cluster map were then imposed onto a pathway diagram to convey the correlations between nodes (metabolites) and edges (reactions) of the graph (metabolic pathway). We adopted a similar approach to analyze our data and visualize the correlated nodes and edges with the goal of formulating better hypotheses about the lipid accumulation pathway. First, a cluster map capturing the correlations between measured transcripts and lipids was constructed for each condition with the horizontal axis representing the mRNA transcripts and the vertical axis consisting of the lipids. Figure S3 shows the cluster map for the correlations between lipids and mRNA transcripts in the baseline condition of C/N 5.

The cluster map in Figure S3 can be partitioned into 4 major quadrants. The reactions and metabolites (lipids) found in each of these quadrants were then assigned a color and these colors were superimposed onto a pathway map depicting the lipid biosynthetic pathways, extracted from a genome-scale model of *R. toruloides* IFO0880 (Dinh et al., 2019). The pathway visualization for lipid accumulation in C/N of 5 is shown in Figure 8 and for C/N of 150 in Figure 9. The Escher maps for visualization were adapted from the genome-scale model developed by Dinh et al (Dinh et al., 2019; King et al., 2015). Reactions displayed using green are correlated positively with storage lipid increase (DAGs and TAGs), whereas those displayed with red are negatively correlated with storage lipid accumulation. The linear chain of reactions upstream of storage lipid synthesis were mostly positively correlated as expected. The reactions of phospholipid synthesis that utilize the same precursors as storage lipids, thus withdrawing flux from storage lipids synthesis are negatively correlated with storage lipid increase. This observation was verified from the lipidomic data where phospholipid values dropped in the oleaginous phase where DAG and TAG increased. The surprising observation, however, was that the synthesis of two phospholipids, phosphatidylinositol (PI) and phosphatidylserine (PS), were positively correlated with storage lipid increase in the C/N 5 (Figure 8).

**Figure 8.**
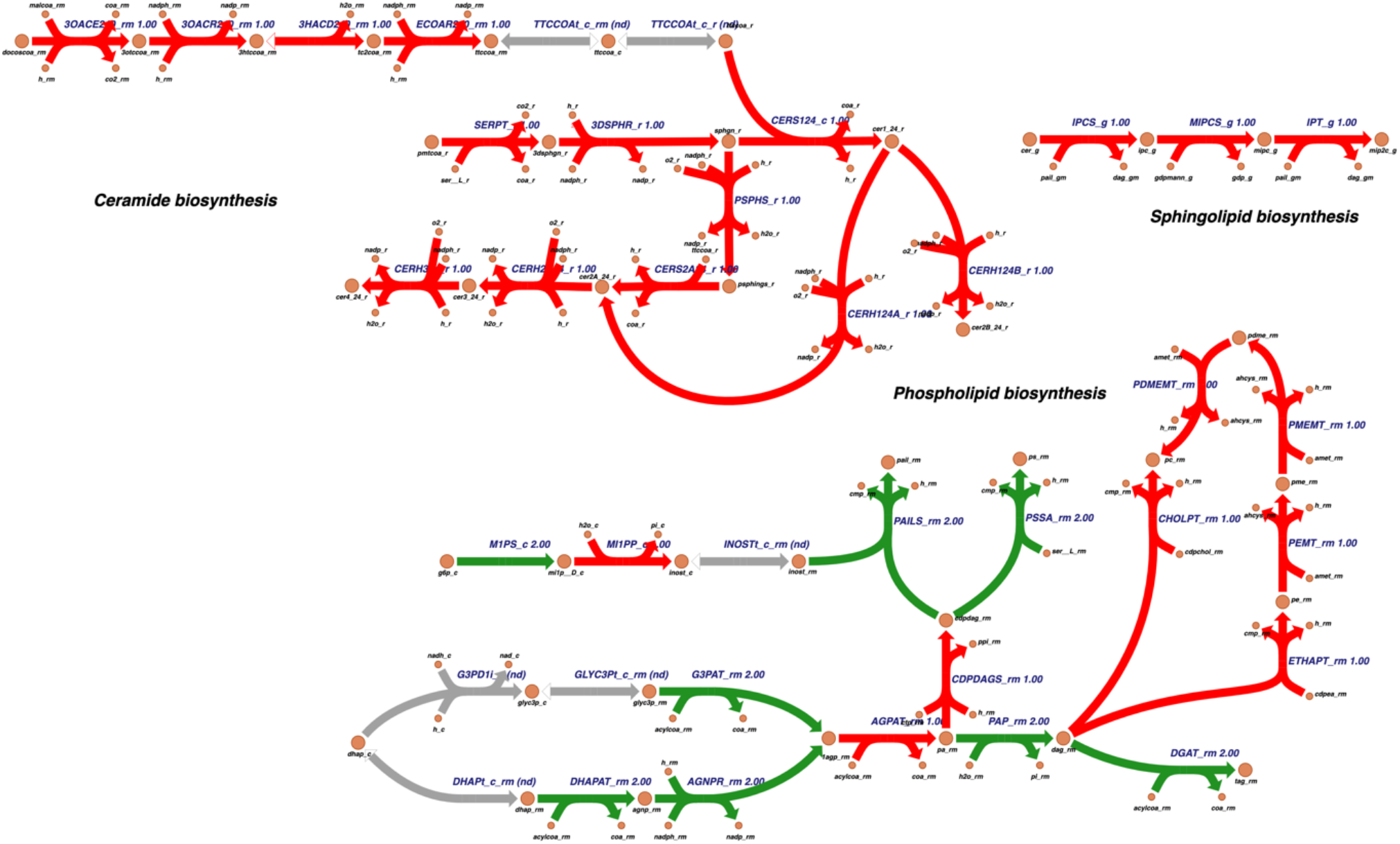
Pathway visualization of the correlation clusters obtained in a cluster map that were then superimposed onto a pathway map, performed for the C/N 5 condition. Reactions in green are positively correlated with storage lipid accumulation (nodes at the bottom right), whereas reactions in red are negatively correlated.

**Figure 9.**
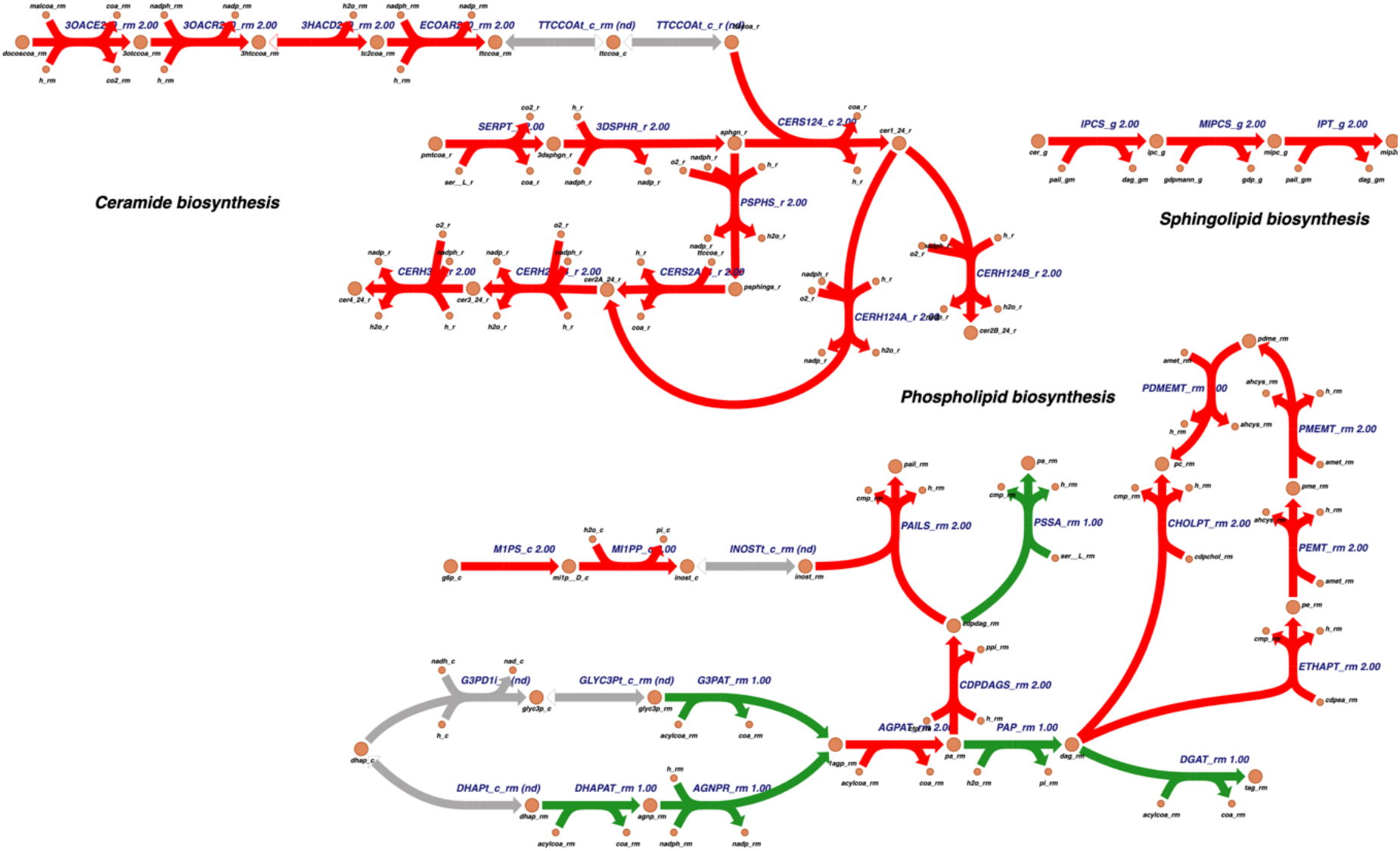
Pathway visualization performed for the C/N 150 culture condition. Reactions in green are positively correlated with an increase in the storage lipids (bottom right nodes), while reactions in red are negatively correlated (based on the clustermap analysis).

The common observation in both Figure 8 and Figure 9 was a similar pattern of selective regulation of reactions within lipid biosynthetic pathways in the context of storage lipid accumulation. The phospholipid biosynthetic pathways were negatively correlated to storage lipid accumulation in both maps (reaction pathway on the right), whereas reactions upstream of the storage lipid synthesis for acylation of the glycerol backbone were upregulated in both (bottom left). These patterns supported the observation that lipid accumulation during nitrogen starvation was accompanied by a rerouting of carbon flux towards storage lipids and away from glycerophospholipids.

## 4. Discussions

The non-model yeast *R. toruloides* is a promising chassis for metabolic engineering due to its ability to accumulate lipids at a much higher fraction than the model yeasts (Chattopadhyay and Maiti, 2021). However, the lipid accumulation phenotype is generally induced under nutrient starvation conditions, where growth is compromised, leading to a tradeoff between lipid levels and biomass (Lopes et al., 2020; Wang et al., 2018). Identifying the mechanism that governs accumulation is thus crucial for any effort to engineer a platform strain that can be employed for overproduction of oleochemicals (Zhu et al., 2012). In this work, we performed a comprehensive study of lipid accumulation phenotypes in *R. toruloides* by integrating transcriptomic and lipidomic analyses to identify the metabolic and regulatory processes that contribute the most to lipid accumulation for engineering purposes. We observed that the transcriptomic profiles for nitrogen-rich and nitrogen-deficient cultures clustered separately, with the divergence of profiles emerging as early as the exponential growth phase. The lipidomic profiles, however, showed greater similarity between the two types of growth conditions. Lipidome trends measured in the C/N of 5 trailed those of C/N 100 or 150 with a time-delay, presumably mirroring the trend of nitrogen consumption. Pathway visualization techniques demonstrated a map of increased regulation within the lipid biosynthesis pathway wherein neighboring reactions were either up- or down-regulated during lipid accumulation. These localized regulations along with trends observed within the lipidomic data proved that the oleaginous yeast *R. toruloides* displayed lipid remodeling during the accumulation phase instead of a uniform increase in all lipids.

Nitrogen starvation was applied in two different carbon-to-nitrogen ratios (C/N of 100 and 150) but both cultures showed similar transcriptomic responses compared to the baseline condition of C/N of 5. Clustering analysis of transcriptomic data showed a clear divergence of trajectories between the baseline and the nitrogen-starved conditions, with the split occurring from the 12-hour timepoint itself (Figure 3). While the divergence of transcriptomic profiles grew more prominent at the saturation phase timepoint of 36 hours, the intracellular lipid profiles showed much less distance in the clustering patterns (Figure 7 and Figure S1). This similarity of lipidomes as the cultures progressed in time can be explained by the fact that, over time, each growth condition (including C/N of 5) would become more nitrogen deficient because nutrients get used up by the cells with nothing being replenished. Further, the intracellular phospholipid profiles of C/N 5 culture at the 36-hour mark resemble the profiles of the C/N 100 and 150 at the earlier 12-hour measurements, suggesting that the C/N 5 condition experiences a similar form of nitrogen starvation at 36 hours of growth as that faced by C/N 100 and 150 at about 12 hours of growth.

The lipidomic timeseries profiles for all 3 conditions followed similar trends (Figure 5) with a drop in mostly phospholipid-based lipidome being accompanied by a rise in mostly storage lipid-based one (DAG and TAG). This trend was demonstrated in Figure 6 where the fraction of phospholipids dropped with increase culture time for all 3 growth conditions, while the fraction of storage lipids increased. Thus, unlike a trend of overall increase in lipids seen in *Y. lipolytica* (Kerkhoven et al., 2016), the increase in lipid pools in *R. toruloides* from nitrogen starvation resulted mainly in the increase of storage lipids. The selective decrease/increase of different lipid pools during the lipid accumulation phase suggested that lipid accumulation is accompanied by strong regulation of the lipid biosynthetic pathways. This contrasts with the findings in *Y. lipoytica* where regulation of the lipid pathways exerted very weak control over the lipid accumulation phenotype (Kerkhoven et al., 2016).

To identify the lipid biosynthetic pathways whose up- or down-regulation most significantly influenced this observed lipid remodeling, the two ‘omic datasets were analyzed in an integrated manner. A cluster map enabled the binning of lipid classes and lipid biosynthetic reactions that showed a similar increase or decrease, and imposing these trends onto a map of the lipid pathways allowed the visualization of these trends with respect to storage lipid levels (Figure 8). Specifically, the pathway visualization showed a clear shutdown of phospholipid biosynthesis pathways under lipid accumulation conditions, which agreed with the decreasing phospholipid levels observed in lipidomic data. More interestingly, the pathway maps also showed that lipid accumulation was accompanied with a positive correlation in the acylglycerol formation from the DHAP pathway, indicating that the increased carbon into the TAG and DAG pools originated from the acylation of the glycerol-3-phosphate backbone.

The multi-omic integrative analysis enabled the identification of reactions that were regulated in a coordinated manner with an increase in storage lipids. The trend of the lipid increase was observed in the direction of storage lipids only rather than a net increase of all intracellular lipid pools. Future studies can leverage the knowledge of these regulated pathways for developing engineering strategies. The strategy of multi-omic analyses for identifying lipid accumulation phenotypes has been employed in other organisms (Jeffers and Roth, 2021). A study by Ajjawi et al. identified a putative set of transcriptional factors (TFs) that governed lipid production in the microalga *Nannochloropsis gaditana* under nitrogen-deficient conditions (Ajjawi et al., 2017). CRISPR-based knockdown of this putative set helped identify one TF that resulted in higher lipid accumulation without affecting growth. Such a method could be adapted to *R. toruloides* using CRISPR-interference systems with the goal of discovering global transcriptional factors that result in a lipid overproducing platform strain.

## 5. Conclusions

In this study, we performed a multi-omic analysis of the lipid accumulation phenotype in the oleaginous yeast *R. toruloides* IFO0880 under nitrogen starvation. Transcriptomic analysis indicated divergent timeseries profiles of the baseline growth media (C/N of 5) compared to the nitrogen-deficient media (C/N of 100 and 150). Lipidomic analysis showed dissimilar lipid profiles at the start with the C/N 5 lipidome eventually converging towards the C/N 100, 150 lipidomes as nitrogen levels decreased over time. Multi-omic analysis of the growth conditions suggested that nitrogen starvation causes the lipidome to remodel with most of the glycerophospholipid pool shifting towards storage lipids. Thus, in batch culture conditions in *R. toruloides*, we observed a selective increase in storage lipids during nitrogen starvation instead of an overall increase of all lipids.

## 6. Acknowledgements

The authors wish to thank Dr. Hoang Dinh and Dr. Patrick Suthers for helpful discussions on metabolic pathway visualization. This work was funded by the U.S. Department of Energy (DE-SC0018260) and the DOE Center for Advanced Bioenergy and Bioproducts Innovation (U.S. Department of Energy, Office of Science, Office of Biological and Environmental Research under Award Number DE-SC0018420). Any opinions, findings, and conclusions or recommendations expressed in this publication are those of the author(s) and do not necessarily reflect the views of the U.S. Department of Energy.

