## Supplementary Figures S1-S3 and Tables S1-S2 for "Nitrogen starvation causes lipid remodeling in *Rhodotorula toruloides*"

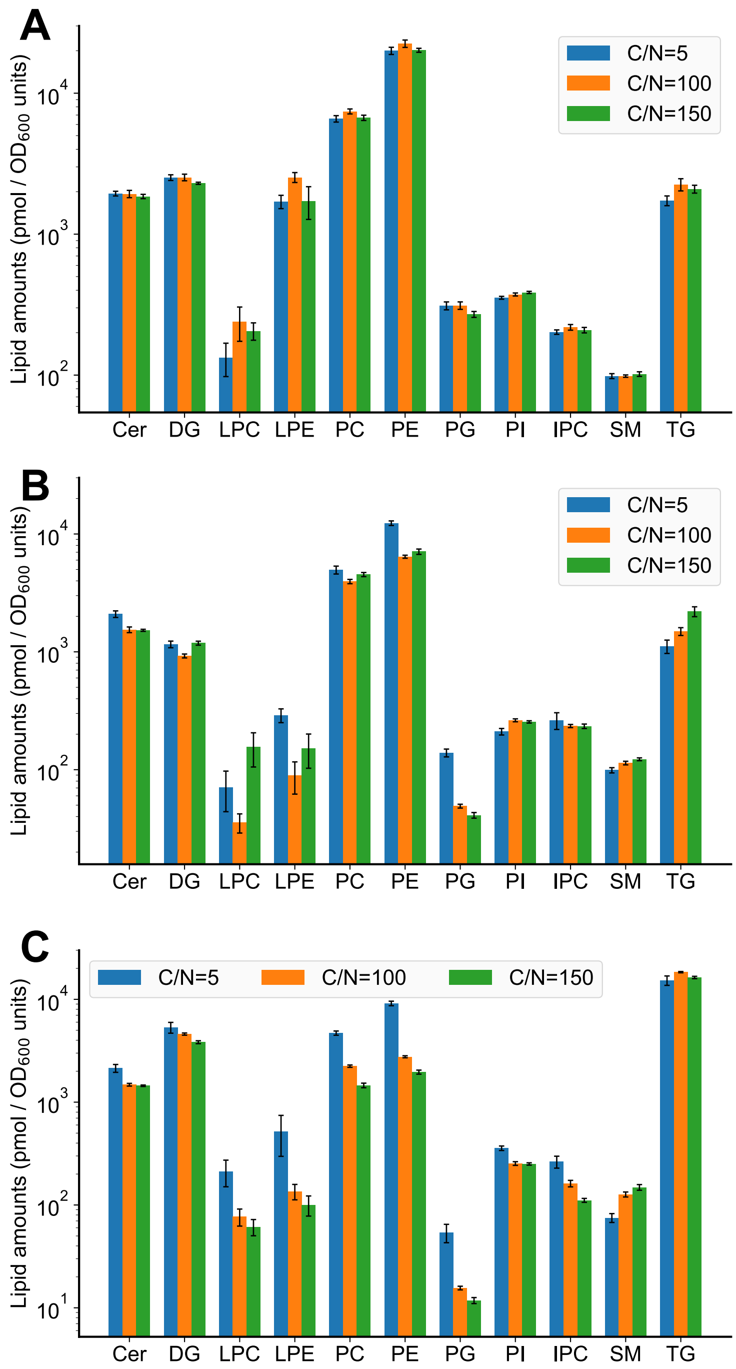


Figure S1. Lipidomic analysis of IFO0880 strain cultivated in different growth media containing C/N ratios of 5, 100 and 150, and sampled at various timepoints. Quantified lipid classes of all three growth conditions sampled after A) 8 hours, B) 12 hours and C) 36 hours of growth.

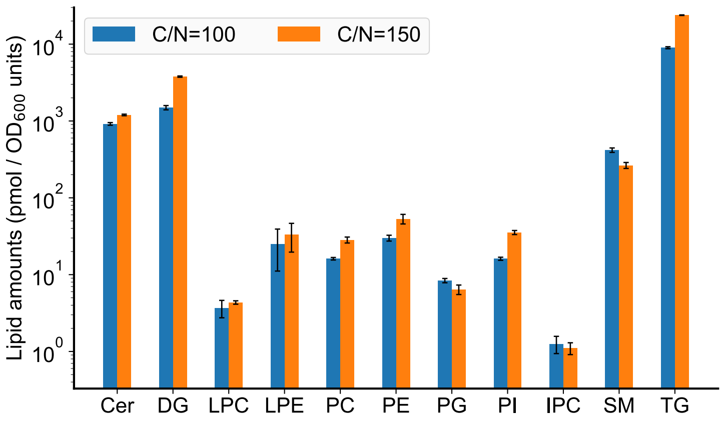


Figure S2. Lipidomic analysis of IFO0880 grown in C/N 100 and 150 culture conditions and sampled at a timepoint of 88 hours (oleaginous phase).

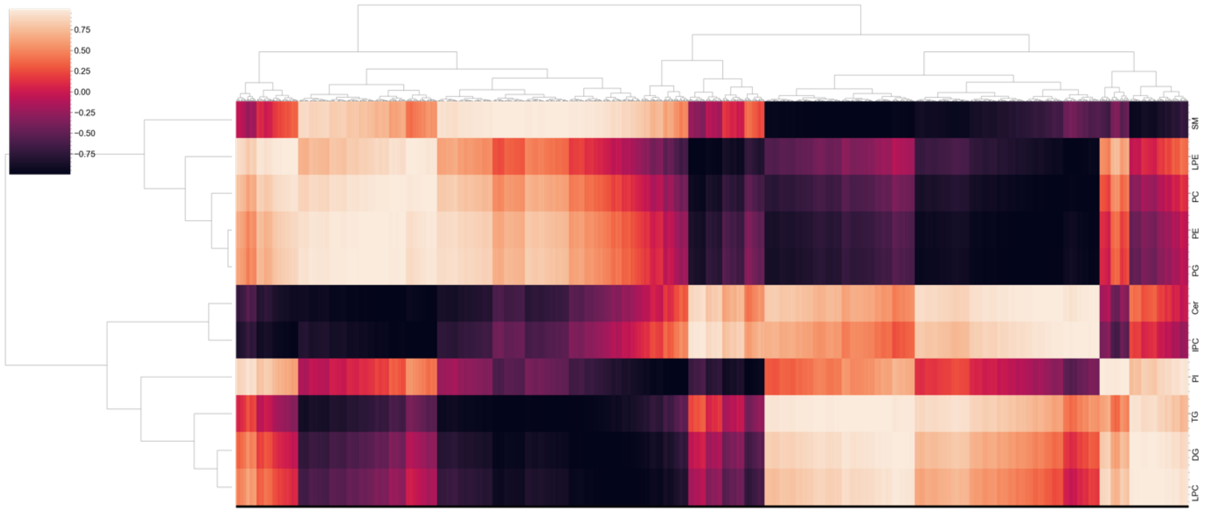

Figure S3: Clustermap for the baseline condition of C/N 5 where the horizontal axis represents the mRNA entries and vertical axis showing the lipid measurements.

Table S1. Media formulations of different C/N ratios used in this study (all amounts listed under C/N 5, 100 and 150 columns are volume amounts in mL to make up a total solution of 100 mL).

| **Stocks** | **Final concentration** | **C/N 5** | **C/N 100** | **C/N 150** |
|---|---|---|---|---|
| Glucose (200 g/L) | 20 g/L | 10 | 10 | 10 |
| YNBwaaas (17 g/L) | 1.7 g/L | 10 | 10 | 10 |
| $NH_4Cl$ (61 g/L) | 6.1 / 0.305 / 0.203 g/L | 10 | 0.5 | 0.33 |
| $Na_2HPO_4$ (0.5M) | 25 mM | 5 | 5 | 5 |
| $KH_2PO_4$ (1M) | 125 mM (pH = 5.6) | 12.5 | 12.5 | 12.5 |
| Water | | 52.5 | 62 | 62.17 |

*YNBwaaas: Yeast Nitrogen Base without amino acids or ammonium sulphate

Table S2. Composition of internal standards spiked-in during lipidomic extraction.

| Class | Component | Amount (pmol) |
|---|---|---|
| **PC** | IS PC 15:0/18:1-d7 | 200.10 |
| **PE** | IS PE 17:0/14:1 | 14.58 |
| **PS** | IS PS 17:0/14:1 | 13.44 |
| **PG** | IS PG 15:0/18:1-d7 | 34.96 |
| **PI** | IS PI 17:0/14:1 | 12.59 |
| **PA** | IS PA 15:0/18:1-d7 | 10 |
| **LPC** | IS LPC 18:1-d7 | 45 |
| **LPE** | IS LPE 18:1-d7 | 10.08 |
| **EE** | IS CE 18:1-d7 | 500.40 |
| **DAG** | IS DAG 15:0/18:1-d7 | 14.98 |
| **TAG** | IS TAG 15:0/18:1-d7/15:0 | 65.04 |
| **Cer** | IS Cer 18:1;2/17:0 | 226.48 |
| **IPC** | IS PI 17:0/14:1 | 12.59 |
| **MIPC** | IS PI 17:0/14:1 | 12.59 |
| **MIP2C** | IS PI 17:0/14:1 | 12.59 |
| **Ergosterol** | IS Cholesterol-d7 | 250.12 |
| **LPA** | IS PA 15:0/18:1-d7 | 10 |
| **LPI** | IS PI 17:0/14:1 | 12.59 |
| **LPS** | IS PS 17:0/14:1 | 20.16 |
